# HybridMolDB: a manually curated database dedicated to hybrid molecules for chemical biology and drug discovery

**DOI:** 10.1101/516765

**Authors:** Yecheng Li, Chongze Zhao, Boyan Wei, Ling Wang

**Affiliations:** Joint International Research Laboratory of Synthetic Biology and Medicine, School of Biology and Biological Engineering, South China University of Technology, Guangzhou 510006, China

## Abstract

**Summary:** Hybrid molecule-based drug design is the combination of two or more pharmacophoric moieties from identical or non-identical bioactive molecules, or known identical or non-identical bioactive molecules, into a single chemical entity. This strategy may be used to achieve better affinity and efficacy, or improved properties compared to the parent molecules, to interact with two or multiple targets, to reduce undesirable side effects, to decrease drug-drug interactions or to reduce emergence of drug resistance. The approach offers the prospect of better drugs for the treatment of many human diseases. Research activity in this area is increasing and has attracted many practitioners worldwide. To accelerate the design and discovery of new hybrid molecule-based drugs, it is essential to properly collect and annotate experimental data obtained from known hybrid molecules. To address this need, we have developed HybridMolDB, a manually curated database dedicated to hybrid molecules for chemical biology and drug discovery. It contains structures, manually annotated design protocols, pharmacological data, some physicochemical, ligand efficiency, drug-likeness and ADMET characteristics, and the biological targets of known hybrid molecules. HybridMolDB supports a range of query types, including searches by extensive text, protein sequence, chemical structure similarity and property ranges.

**Availability:** HybridMolDB is freely available at http://www.idruglab.com/HybridMolDB/index.php.

## 1 INTRODUCTION

Hybrid molecule-based drug discovery is an appealing and emerging discipline in medicinal chemistry. The strategy is to design new molecules by hybridizing two or more pharmacophoric moieties from known identical or non-identical bioactive molecules, or two or more known identical or non-identical bioactive molecules, into a single molecule (Viegas-Junior, et al., 2007). Hybrid molecules can be divided into two major categories: linked hybrid molecules (LHM) and fused hybrid molecules (FHM). The LHM category contains two subcategories: direct (DLHM) and spacer (SLHM).

Hybrid molecules have been shown to be useful for generating better and specific affinity and efficacy, or interacting with two or multiple targets, or reducing undesirable side effects, or diminishing the likelihood of drug resistance, as well as enhancing patient compliance (Abdolmaleki and Ghasemi, 2017). Given the advantages of hybrid molecule-based drug discovery, both industrial and academic practitioners have designed such molecules for the treatment of many human diseases (Chugunova and Burilov, 2017; Fortin and Berube, 2013; Korth, et al., 2013). Hybrid molecules can be attractive drug candidates, but there is no specific resource that provides comprehensive experimental data on hybrid molecules to support such therapeutic innovation.

Herein, we have developed a hybrid molecule database (HybridMolDB) to systematically collect, validate and annotate known hybrid molecules from publicly available resources.

## 2 METHOD

### 2.1 Data collection and processing

The PubMed and ChEMBL_23 (Bento, et al., 2014) databases were searched for scientific publications on hybrid molecules by using combinations of the following keywords: ‘hybrid’, ‘conjugate’, ‘fusion’, ‘fused’, ‘mixed’, ‘merge’, ‘blend’, ‘combinate’, ‘combination’, ‘bear’, ‘link’, ‘spacer’, ‘dimer’, ‘dual’, ‘bifunctional’, ‘multifunctional’, ‘bivalent’ and ‘bitopic’. The search results were further refined, based on the definition of hybrid molecule, by manual inspection. Comprehensive information on the hybrid molecules was mined manually from each peer-reviewed research article. The chemical structural information for the hybrid molecule and the parent bioactive molecules were derived from ChEMBL. For those hybrid molecules and/or precursor ligands in the respective literature that were not found in ChEMBL, their structures were hand drawn using Marvin Sketch software. All two- and three-dimensional structures of the identified hybrid molecules in multiple formats (SMILES, MOL, MOL2, SDF, PDB and PDBQT) were uniformly converted using OpenBabel (O’Boyle, et al., 2011). The design protocol for each hybrid molecule was manually annotated from the original literature. The pharmacological data on the hybrid molecule was retrieved from ChEMBL or was hand-picked from the original literature. Associated data, including biological targets (genes, functions, sequences, structures, signal transduction pathways and diseases etc.) and target related approval or experimental drugs, were retrieved from UniProt (Bateman, et al., 2015), TTD (Yang, et al., 2016) and DrugBank (Law, et al., 2014) databases. Finally, a series of ligand properties (Part I of the Supplementary Material and Table S1) were calculated for each hybrid molecule.

### 2.2 Database and web interface implementation

HybridMolDB was implemented as a relational database in MySQL. The web interface was built using HTML, PHP and JavaScript. Four retrieval methods (text, chemical structure, protein sequence, and physicochemical property threshold criteria searches, Part I of the Supplementary Material) are provided to query HybridMolDB.

## 3 Results and discussion

### 3.1 Database content and statistics

HybridMolDB contains 4927 unique, experimentally validated hybrid molecules. The proportions of LHM and FHM in the database are 47.23% and 52.77% (Fig. S1A). Approximately 46.21% of the LHM are SLHM that include 45.26% non-cleavable and 0.95% cleavable SLHM. Only 1.01% of LHM are DLHM that are cleavable hybrid molecules. A total of 4927 annotated design protocols and 35,363 pharmacological data points for the hybrid molecules are stored in HybridMolDB. These pharmacological data (Fig. S1B) are from target-, cell-, and organism-based assays, comprising 15.83%, 31.56% and 33.12%, respectively. A total of 260 target diseases are associated with the collected hybrid molecules and the top nine are shown in Fig. S2A. Moreover, a total of 1830 annotated hybrid molecules are active against 107 biological targets, and approximately 62.74% of these hybrid molecules act on two or more targets (Fig. S2B). A total of 107 biological targets and related information (genes, functions, sequences, structures, signal transduction pathways, related diseases and drugs, etc.) associated with these hybrid molecules were collected and stored in the database.

### 3.2 Web interface and usage

A detailed description of the web interface and its usage are described in Part II of the Supplementary Material and Figs. S3 and S4.

## 4 CONCLUSION

Given the importance of hybrid molecules in the field of chemical biology and drug discovery, we have developed HybridMolDB that systematically collects and annotates known hybrid molecules from published scientific literature. It is hoped that the data in HybridMolDB and the pre-calculated ligand properties of known hybrid molecules, with freely accessible and versatile tools to query them, will accelerate drug discovery and chemical biology projects for such molecules. In addition, big data analysis of experimentally-validated hybrid molecule data, together with chemical structures and properties of known hybrid molecules in the database, should facilitate development and/or optimization of related *in-silico* tools.

## Funding

This work was supported in part by the Science and Technology Program of Guangzhou (no. 201707010063), the National Natural Science Foundation of China (no. 81502984), the Natural Science Foundation of Guangdong Province (no. 2016A030310421), and the Medical Scientific Research Foundation of Guangdong Province (no. A2018114).

### Conflicts of Interest

none declared.

## References

Abdolmaleki, A. and Ghasemi, J.B. (2017) Dual-acting of hybrid compounds - a new dawn in the discovery of multi-target drugs: lead generation approaches. Curr. Top. Med. Chem., 17, 1096–1114.

Bateman, A., et al. (2015) UniProt: a hub for protein information. Nucleic Acids Res., 43, D204–D212.

Bento, A.P., et al. (2014) The ChEMBL bioactivity database: an update. Nucleic Acids Res., 42, D1083–D1090.

Chugunova, E.A. and Burilov, A.R. (2017) Novel structural hybrids on the base of benzofuroxans and furoxans. mini-review. Curr. Top. Med. Chem., 17, 986–1005.

Fortin, S. and Berube, G. (2013) Advances in the development of hybrid anticancer drugs. Expert Opin. Drug Discov., 8, 1029–1047.

Korth, C., Klingenstein, R. and Muller-Schiffmann, A. (2013) Hybrid molecules synergistically acting against protein aggregation diseases. Curr. Top. Med. Chem., 13, 2484–2490.

Law, V., et al. (2014) DrugBank 4.0: shedding new light on drug metabolism. Nucleic Acids Res., 42, D1091–D1097.

O’Boyle, N.M., et al. (2011) Open Babel: An open chemical toolbox. J. Cheminf., 3, 33.

Viegas-Junior, C., et al. (2007) Molecular hybridization: a useful tool in the design of new drug prototypes. Curr. Med. Chem., 14, 1829–1852.

Yang, H., et al. (2016) Therapeutic target database update 2016: enriched resource for bench to clinical drug target and targeted pathway information. Nucleic Acids Res., 44, D1069–D1074.

