## Supplemental Figure1-4 Table 1 for "HybridMolDB: a manually curated database dedicated to hybrid molecules for chemical biology and drug discovery"

### Part I: Search tools and ligand properties calculation

HybridMolDB provides four methods to retrieve hybrid molecules and their associated data: text mining, chemical structure search, protein sequence search, and physicochemical property thresholds criteria search. Two algorithms for searching by chemical structure are employed in HybridMolDB: substructure search and two-dimensional (2D) similarity calculation. The 2D similarity calculations are based on the FP2 fingerprint and performed using OpenBabel (O'Boyle, et al., 2011). The similarity method uses the Tanimoto coefficient to quantify similarity between two molecules. The BLAST algorithm is used for protein sequence similarity searching (Altschul, et al., 1990).

A series of ligand properties (if computable, Supplementary Table S1), including 17 physicochemical, nine drug-likeness, seven ligand efficiency metrics and 26 ADMET (absorption, distribution, metabolism, excretion and toxicity) characteristics were calculated for each hybrid molecule. The physicochemical and drug-likeness properties were calculated using FAF-Drugs4 (Lagorce, et al., 2017) and Discovery Studio software package (v3.5; Biovia, San Diego, CA, USA). Ligand efficiency metrics (Hopkins, et al., 2014) based on biological target activity values (e.g., $IC_{50}$, $K_i$, $K_d$, and $EC_{50}$, if available) were computed using an *in-house* script. The ADMET characteristics were generated using admetSAR (Cheng, et al., 2012).

These ligand properties are commonly used to evaluate physicochemical, lead- and drug-likeness, and pharmacokinetics profiles of hybrid molecules. The current version of HybridMolDB contains 83616 physicochemical property data points, 41163 druggability characteristics data points, 22029 ligand efficiency metrics values, and 128102 ADMET data points (Supplementary Table S1). Lipinski's rule of five (Lipinski, et al., 1997), Veber rule (Veber, et al., 2002) and ADMET properties for each hybrid molecule are normalized statistics and rendered in online radar charts using Chart.js (http://www.chartjs.org/), which enables users to intuitively determine the quality of a given property.

**Part II: Web interface and usage**

HybridMolDB provides a user-friendly web interface for users to search, browse, display, and download all experimentally validated hybrid molecule data in the database. Moreover, public users and practitioners can send new hybrid molecule data to us through the Contribute Data module encoded in HybridMolDB.

***Search.*** HybridMolDB provides four modes to query the database: text search (target name, UniProt ID, SMILES, ChEMBL ID and InChIKey), substructure and 2D structural similarity, protein sequence similarity to HybridMolDB target entries, and physicochemical properties (e.g., molecular weight, hydrophobicity, polar surface area) thresholds criteria search. Herein, an example 2D similarity search is provided to show the use of the HybridMolDB search function (Supplementary Fig. S3A). We sought to capture information on fimepinostat (CUDC-907, a dual HDAC/PI3K inhibitor) that was designed by merging pharmacophoric moieties from pictilisib (PI3K inhibitor) and quisinostat (HDAC inhibitor) (Chen, et al., 2018). Fimepinostat is being investigated in a phase 2 clinical trial for the treatment of relapsed/refractory lymphomas or multiple myeloma and in a phase 1 trial in patients with advanced solid tumors (Younes, et al., 2016).

The user can either copy and paste a SMILES string or directly sketch the molecular structure of fimepinostat within the Marvin JS editor (https://chemaxon.com/). Clicking "Fetch Compounds" performs a 2D structural similarity search against the hybrid molecules in HybridMolDB. The 2D similarity search results are ranked by their Tanimoto similarity score and are shown in Supplementary Fig. S3B. The first record is for fimepinostat, which has the highest similarity score value of 1. The user can then click the "Show Detail" button to view the hybrid molecule ID card of fimepinostat (Supplementary Fig. S3C). The compound card displays all of the data on fimepinostat via different tabs: basic information (structure, component precursors, compound ID, molecular formula and classification), annotated design protocol, identifiers (InChIKey and canonical SMILES), pharmacological activities (biological assay related target name and type, activity value, description, PubMed ID, etc.), ligand properties (physicochemical, drug-likeness, ligand efficiency metrics, and ADMET characteristics), property statistics based on Lipinski's rule of five, Veber rule and ADMET using online radar charts and downloads (Supplementary Fig. S3C). In the pharmacological activities table, the user can click the "Target Name" hyperlink (e.g., histone deacetylase, Supplementary Fig. S3C) to check and browse detailed target related information (Supplementary Fig. S3D) including gene, function, sequences, structures, signal transduction pathways, related diseases, and drugs. In addition, all 2D and 3D structures of fimepinostat in multiple formats (SMILES, MOL, MOL2, SDF, PDB and PDBQT) can be downloaded (Supplementary Fig. S3C).

***Browse.*** Hybrid molecules can be browsed by their classifications (LHM and FHM) in the "Browse" page. Taking LHM as an example hybrid molecule classification (Supplementary Fig. S4), according to linker bond type (e.g., –C–C–, –C–N, –C–O–), users can directly browse the desired hybrid molecule data. In addition, the Search module encoded in the "Browse" page enables users to fetch and browse the desired hybrid molecule by inputting its InChIKey.

***Download & FAQ.*** All metadata in HybridMolDB can be freely downloaded from the "Download" page. A detailed introduction to HybridMolDB and a tutorial are available on the "FAQ" page.

***Contribute Data.*** If public users and practitioners know or have new experimentally-validated hybrid molecule data that they would like added, the PubMed ID or Digital Object Identifier (DOI) can be submitted and/or scientific articles in PDF format can be uploaded directly via the 'Contribute Data' page. The information is sent to us using the Submit module on the "Contribute Data" page and we will process the received hybrid molecule information and add it to the database.

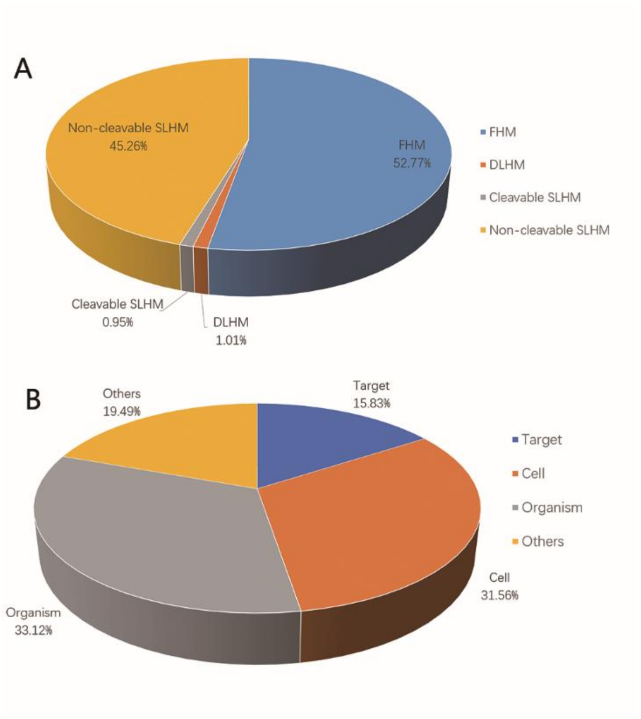


**Supplementary Fig. S1.** Distribution of different hybrid molecule categories (**A**) and pharmacological data for all collected hybrid molecules (**B**) in HybridMolDB. FHM: fused hybrid molecules; LHM: linked hybrid molecules; LHM contains two subcategories: direct and spacer LHM (DLHM and SLHM). SLHM can be further divided into two subclasses: cleavable or non-cleavable SLHM.

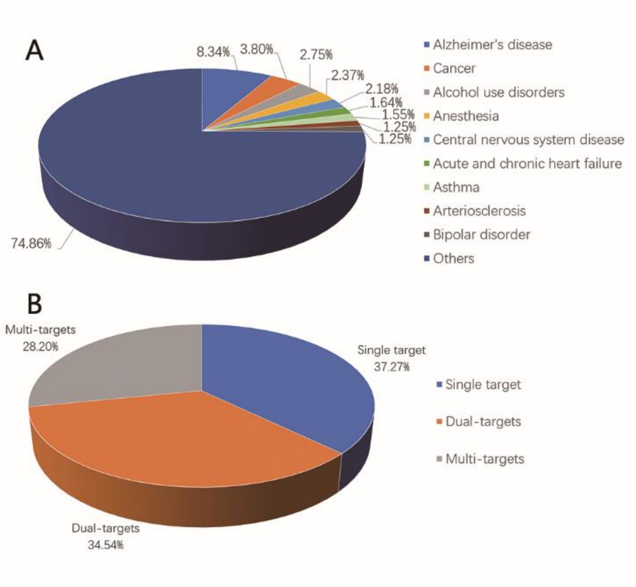


**Supplementary Fig. S2.** Top nine diseases associated with all collected hybrid molecules **(A)** and distribution of binding modes for 1830 hybrid molecules from target-based pharmacological data **(B)**.

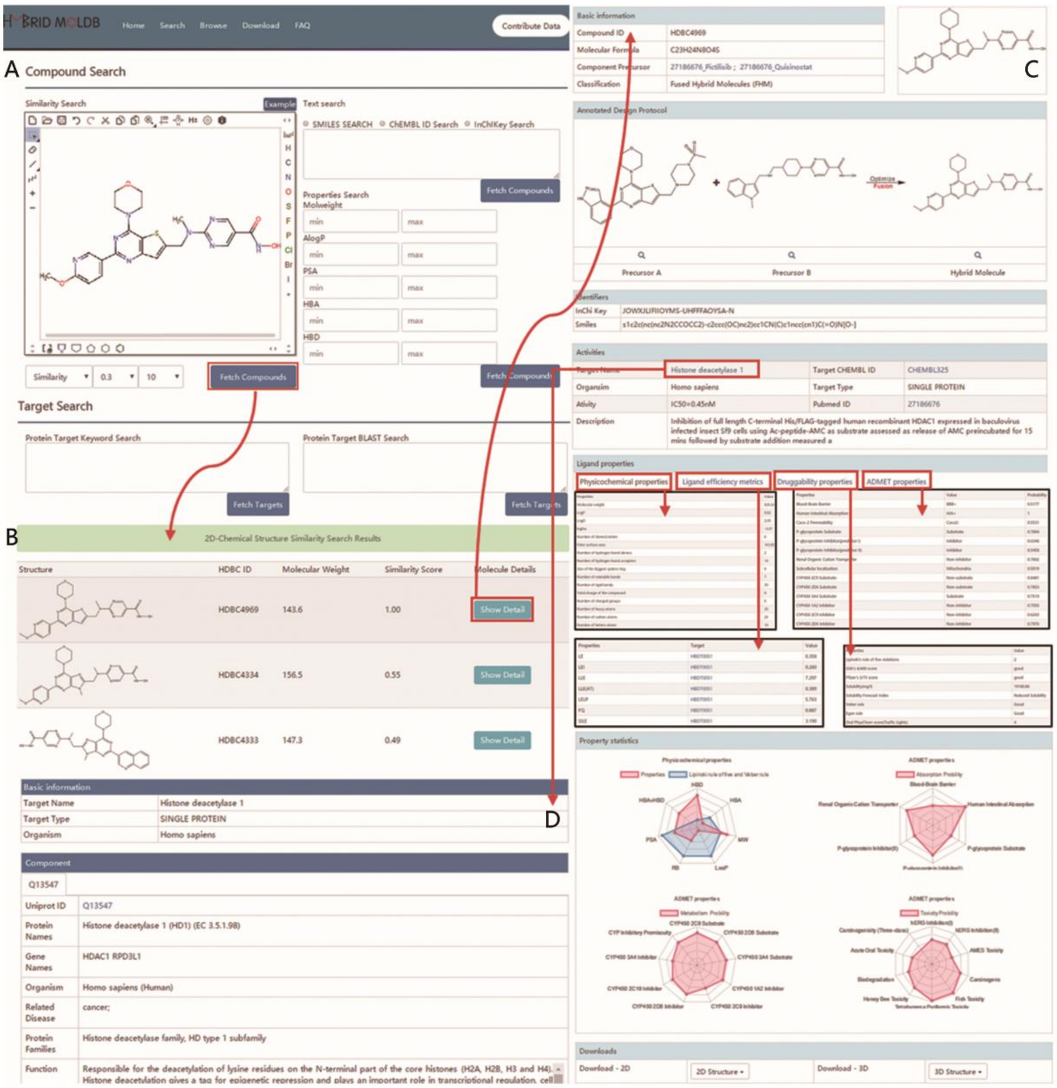


**Supplementary Fig. S3.** A schematic workflow of the chemical structure search and results display interface in HybridMolDB. (**A**) 2D chemical similarity for fimepinostat drawn using the online Marvin JS sketcher. (**B**) Snapshot of search results for fimepinostat obtained from the 2D similarity search. (**C**) Snapshot of the hybrid molecule ID card of fimepinostat. (**D**) Detailed target related information for histone deacetylase.

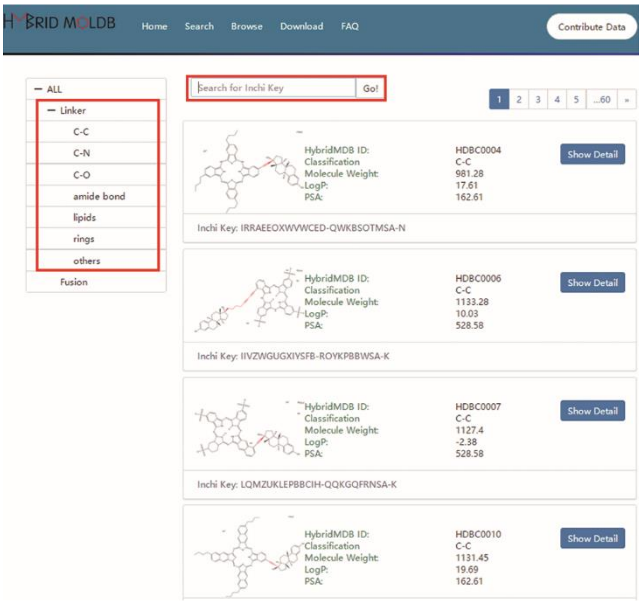


**Supplementary Fig. S4.** A snapshot of the “Browse” function in HybridMolDB.

**Supplementary Table S1**. The detailed ligand properties calculated for each hybrid molecule in HybridMolDB.

| Physicochemical | Drug-likeness | Ligand efficiency metrics | ADMET characteristics | No. of data points |
|---|---|---|---|---|
| Molecular weight<br>LogP<br>logSw<br>Number of stereocenters<br>Polar surface area<br>Number of hydrogen bond donors<br>Number of hydrogen bond acceptors<br>Size of the biggest system ring<br>Number of rotatable bonds<br>Number of rigid bonds<br>Total charge<br>Number of charged groups<br>Number of heavy atoms<br>Number of carbon atoms<br>Number of hetero atoms | Lipinski's rule of five violations<br>GSK's 4/400 score<br>Pfizer's 3/75 score<br>Solubility (mg/L)<br>Solubility forecast index<br>Veber rule<br>Egan rule<br>Oral PhysChem score(Traffic Lights)<br>Quantitative estimate of drug-likeness | LE (Ligand efficiency)<br>LEI (Ligand efficiency index)<br>LLE (Lipophilic ligand efficiency)<br>LLE(AT) ( LLE adjusted for heavy atom count)<br>LELP (Lipophilicity-corrected ligand efficiency)<br>FQ (Fit quality)<br>SILE (Size-independent ligand efficiency) | Blood-brain barrier<br>Human intestinal absorption<br>Caco-2 permeability<br>P-glycoprotein substrate<br>P-glycoprotein inhibitor (predictor I)<br>P-glycoprotein inhibitor (predictor II)<br>Renal organic cation transporter<br>Subcellular localization<br>CYP450 2C9 substrate<br>CYP450 2D6 substrate<br>CYP450 3A4 substrate<br>CYP450 1A2 inhibitor<br>CYP450 2C9 inhibitor<br>CYP450 2D6 inhibitor<br>CYP450 2C19 inhibitor<br>CYP450 3A4 inhibitor<br>CYP inhibitory promiscuity<br>Human Ether-a-go-go-Related Gene inhibition (predictor I)<br>Human Ether-a-go-go-Related Gene inhibition (predictor II)<br>AMES Toxicity<br>Carcinogens<br>Fish Toxicity<br>Tetrahymena Pyriformis Toxicity<br>Honey Bee Toxicity<br>Acute Oral Toxicity<br>Carcinogenicity (Three-class) | Physicochemical properties (83613)<br>Druggability characteristics (41163)<br>Ligand efficiency metrics values (22029)<br>ADMET (128102) |

The ligand efficiency metrics are calculated as follows:

$LE = (1.37/HA) \times pIC_{50}$;

$LEI = pIC_{50}/HA$;

$LLE = pIC_{50} - cLogP$;

$LLE(AT) = 0.111 + 1.37(LLE/HA)$;

$LELP = cLogP/LE$;

$FQ = [pIC_{50}/HA]/[0.0715 + (7.5328/HA) + (25.7079/HA^2) - (361.4722/HA^3)]$, $SILE = pIC_{50}/HA^{0.3}$).

HA: Number of heavy atoms

### References


Altschul, S.F., *et al.* (1990) Basic local alignment search tool. *J. Mol. Biol.*, **215**, 403–410.

Chen, D., *et al.* (2018) Design, Synthesis, and Preclinical Evaluation of Fused Pyrimidine-Based Hydroxamates for the Treatment of Hepatocellular Carcinoma. *J. Med. Chem.*, **61**, 1552–1575.

Cheng, F., *et al.* (2012) admetSAR: a comprehensive source and free tool for assessment of chemical ADMET properties. *J. Chem. Inf. Model.*, **52**, 3099–3105.

Hopkins, A.L., *et al.* (2014) The role of ligand efficiency metrics in drug discovery. *Nat. Rev. Drug Discov.*, **13**, 105–121.

Lagorce, D., *et al.* (2017) FAF-Drugs4: free ADME-tox filtering computations for chemical biology and early stages drug discovery. *Bioinformatics*, **33**, 3658–3660.

Lipinski, C.A., *et al.* (1997) Experimental and computational approaches to estimate solubility and permeability in drug discovery and development settings. *Adv. Drug Deliv. Rev.*, **23**(1–3), 3–25.

O'Boyle, N.M., *et al.* (2011) Open Babel: An open chemical toolbox. *J. Cheminf.*, **3**, 33.

Veber, D.F., *et al.* (2002) Molecular properties that influence the oral bioavailability of drug candidates. *J. Med. Chem.*, **45**, 2615–2623.

Younes, A., *et al.* (2016) Safety, tolerability, and preliminary activity of CUDC-907, a first-in-class, oral, dual inhibitor of HDAC and PI3K, in patients with relapsed or refractory lymphoma or multiple myeloma: an open-label, dose-escalation, phase 1 trial. *Lancet Oncol.*, **17**, 622–631.
